# Zebrafish Foxl2l functions in proliferating germ cells for female meiotic entry

**DOI:** 10.1101/2024.07.30.605757

**Authors:** Ching-Hsin Yang, Yan-wei Wang, Chen-wei Hsu, You-Jiun Pan, Bon-chu Chung

## Abstract

Zebrafish sex differentiation is a complicated process and the detailed mechanism has not been fully understood. Here we characterized a transcription factor, Foxl2l, that participates in female oogenesis. We show that it is expressed specifically in proliferating germ cells in juvenile gonads and mature ovaries. We have used CRISPR-Cas9 to generate zebrafish deficient in *foxl2l* expression. Zebrafish with *foxl2l^-/-^* are all males, and this female-to-male sex reversal cannot be reversed by *tp53* mutation, indicating this sex reversal is unrelated to cell death. We have generated transgenic fish expressing GFP under the control of *foxl2l* promoter to track the development of *foxl2l^+^*-germ cells, which failed to enter meiosis and were accumulated as cystic cells. Our RNA-seq analysis also showed the reduced expression of genes in meiosis and oogenesis among other affected pathways. All together, we show that zebrafish Foxl2l is a nuclear factor controlling the expression of meiotic and oogenic genes, and its deficiency leads to defective meiotic entry and the accumulation of premeiotic germ cells.

**Highlights:**

1. Zebrafish *foxl2l* is expressed only in proliferating germ cells in juvenile gonads and mature ovaries.
2. Foxl2l is a nuclear factor that promotes expression of genes involved in meiosis and oogenesis.
3. Zebrafish depleted of *foxl2l* lack meiotic oocytes in juveniles and become all males in adults.
4. Mutation of *foxl2l* leads to the accumulation of premeiotic germ cells.

## Introduction

### Zebrafish female gonads develop before males

Zebrafish (*Danio rerio*) is an animal model widely used for research. However, its sex determination mechanism is still unclear. In addition to genes that control sex differentiation, the sex ratio of zebrafish is often affected by environmental factors such as temperature [1], breeding density [2], and feeding [3].

The earliest germ cells in zebrafish, primordial germ cells (PGCs), stay undifferentiated until 10 dpf. After that, PGCs start to differentiate into proliferating mitotic germ cells, some of which enter meiotic prophase I at 14 dpf and go through the stages of leptoene, zygotene, pachytene, diplotene and diakinesis [4]. These oocytes stay arrested at the diplotene I stage until ovulation. Large proportions of perinucleolar oocytes can be observed in zebrafish larvae at 21 dpf, presumably advancing towards female fate. In about half of fish, the oocytes undergo apoptosis at one month of age, and the gonads develop into testes [5]. In some fish, mitotic germ cells do not enter meiosis at 14 to 21 dpf, but directly differentiate into spermatocytes at around one month of age [6].

### Known key genes

The domesticated zebrafish strains do not have the sex chromosome and their sex differentiation can be affected by many genes [3]. Some genes affect female cell fate while others affect female meiosis. *Factor in the germline a* (*figla*), which encodes an oocyte-specific transcription factor, is required for oocyte formation; its deficiency results in an all-male phenotype [7]. Meiotic genes, such as *meiosis specific with OB-fold* (*meiob*), *DNA meiotic recombinase 1* (*dmc1*) and *synaptonemal complex protein 3* (*sycp3*), are upregulated in the germ cells in presumptive females at 14 dpf [8]. Depletion of such meiotic genes as *sycp3* [8] and *RAD21 Cohesin Complex Component Like 1* (*rad21l1*) [9] all lead to the development of 100% males. Estrogen production is also important, as the depletion of the gene involved in its synthesis, *cytochrome P450 family 19 subfamily A member 1* (*cyp19a1*), leads an all-male phenotype in adults [10].

Forkhead-box protein L2 (Foxl2) is a conserved ovary-specific factor which regulates oocyte formation and prevents ovary from differentiating into testis [11]. In females, Foxl2 stimulates steroid synthesis by activate *cyp19a* expression in Japanese flounder [12]. Foxl2 depletion yields complete female-to-male sex reversal in goat and Nile tilapia [13, 14]. Foxl2 expression in the mammalian testis is repressed by Doublesex and mab-3 related transcription factor 1 (Dmrt1) to ensure normal testis development [15]. In zebrafish, there are two *foxl2* duplicates, *foxl2a* and *foxl2b* [11]. The gonad of *foxl2a* mutant only contains degenerating oocytes and somatic cells in ovary [16]. The *foxl2b* female mutants exhibit partial sex reversal with gonadal structure similar with testis [16]. Foxl2a and Foxl2b cooperatively regulate ovary maintenance in zebrafish.

### Role of foxl2l in medaka

In medaka (*Oryzias latipes*), a *foxl2* paralogue, *forkhead-box protein L3* (*foxl3*), is expressed in cyst-forming germ cells before germ cell initiates meiosis at bipotential stage, but maintained only in female mitotic germ cells after gonad dimorphism [17]. In *foxl3* depleted medaka, both oocytes and functional sperms are produced in the adult XX ovaries, indicating the role of *foxl3* in suppressing spermatogenesis [17]. *REC8 meiotic recombination protein a* (*rec8a*) and *F-box protein 47* (*fbxo47*) are the direct targets of *foxl3* to activate two independent pathways of mitosis and oogenesis to switch germlines into oocytes [18].

Foxl3 in medaka has been well documented, but the understanding of the zebrafish orthologue, *forkhead-box protein L2-like* (*foxl2l*), is relatively poor. Recent single cell RNA-seq data found that *foxl2l* is highly expressed in mitotic germ cells and co-expressed with *rec8a* in zebrafish indicating *foxl2l in* zebrafish might have the similar functions of *foxl3* in medaka [19]. The heterozygous GFP knock-in mutant zebrafish has normal sex ratio, but all homozygous knock-in mutants are fertile males indicating that *foxl2l* is essential for female development in zebrafish [19]. However, the expression pattern and function of *foxl2l* are still unclear. Therefore, the aims of the current study are investigating the expression pattern and function of *foxl2l* in zebrafish gonad.

## Materials and Methods

### Fish

The zebrafish (*Danio rerio*) strain used in this study is *Tupfel long fin* (*TL*). The protocols for the care and treatment of zebrafish were approved by the Institutional Animal Care and Use Committee of Academia Sinica (IACUC: 10-10-084).

Transgenic zebrafish were generated as described previously [20]. 25 pg of pT2KXIG-*foxl2l*:EGFP plasmid and 25 pg transposase mRNA were co-injected into one-cell staged embryo to generate transgenic fish.

The CRISPR/Cas9 method was used to generate the *foxl2l* mutant zebrafish. gRNA target sites of *foxl2l* were designed by CHOPCHOP and CRISPRscan. Two gRNAs (each 50 pg) and 200 pg of TrueCut Cas9 Protein v2 (Life Technologies, Cat. A36498) were co-injected into embryos before the two-cell stage. Adult F0 founders were sacrificed for gonadal histology analysis after mating with wild type fish to get heterozygous F1 offspring as described previously [21]. Samples were stained with Hematoxylin/Eosin (HE) according to the standard protocol.

The *tp53^zdf1^* mutant zebrafish was obtained from Zebrafish International Resource Center [22].

### Whole-mount *in situ* hybridization and immunohistochemistry

RNA probe was produced from plasmid or PCR product template by *in vitro* transcription using Digoxigenin/Fluorescein (Roche, Cat. 11277073910/11685619910) uridine-5’-triphosphate (UTP) and T7 /SP6 RNA polymerase (Roche, Cat. 10881775001/10810274001).

Fish samples were fixed in 4% paraformaldehyde (PFA) overnight at 4°C. For whole mount *in situ* hybridization, samples were then dehydrated by methanol before probe hybridization. *In situ* hybridization was performed as previously [4, 23]. Briefly, rehydrated samples were digested with Proteinase K (10 μg/mL) and incubated with RNA probes (1 ng/μL) at 70°C overnight. Then they were incubated with Anti-Digoxigenin alkaline phosphatase antibody (1:5000) (Roche, Cat. 11093274910) overnight at 4°C following by BM Purple (Roche, Cat. 11442074001) staining to visualize the locations of RNA probes in purple color. For fluorescent *in situ* hybridization, samples were incubated Anti-Digoxigenin/Fluorescein peroxidase (1:200) (Roche, Cat. 11207739910/11426346910) for 30 min and stained with Tyramide Signal Amplification Fluorescence Kits (PerkinElmer, Cat. NEL745001KT) and 5 μg/mL 4’,6-Diamidino-2-Phenylindole (DAPI).

For immunofluorescence (IF) staining of EGFP, 4% PFA fixed larval trunks or florescence *in situ* hybridized trunks were washed in PBS with 0.2% Triton X-100 (PBST) and blocked for 1 h at room temperature in 10% normal goat serum. After blocking, samples were placed in blocking solution with anti-GFP antibody (1:200) (Aves Labs, GFP-1020) overnight at 4°C. Trunks were then washed by PBST before incubation with anti-chicken 488 (1:200) (Invitrogen, Cat. A-11039) and 5 μg/mL DAPI in PBST overnight at 4°C. Finally, samples were washed by PBST and whole-mounted in glycerol.

Larvae were incubated with 200 μM 5-ethynyl-2’-deoxyuridine (EdU) for 72 h. After treatment, the fish samples were fixed in 4% PFA overnight at 4°C. Following florescence *in situ* hybridization, EdU was stained by Click-iT® EdU Alexa Fluor® 555 Imaging Kit (Thermo Fisher Scientific, Cat. C10338) according to the protocol of manufacture.

Stained samples were imaged by Z1 upright microscope (Carl Zeiss Inc.) for visible light images or LSM780 (Carl Zeiss Inc.) for confocal images.

### Reverse transcription and RT-PCR

Tissues samples were homogenized in Azol reagent (Arrowtec Life Science, Cat. AzolNC.200) with a hand-held polypropylene homogenizer (Kimble/Kontes) according to manufacturer’s protocol. After RNA extraction, 1 μg RNA was used as a template for reverse transcription using the Maxima Frist Strand cDNA Synthesis kit (Fermentas Int., Cat. K1641) to synthesize cDNA. cDNA was used as a template in PCR reaction and amplified by SuperRed PCR Master Mix (Tools, Cat. TE-SR01).

### Cell sorting and RNA-seq

Wildtype or *foxl2l*^-/-^ zebrafish (21 dpf) containing *foxl2l:EGFP* transgene were used for RNA-seq (4-10 fish/sample, triplicates in each genotype). To isolate GFP^+^ germ cell, head, tail and intestine were removed from the trunk and cells in the remaining trunk were dissociated by 0.2% collagenase (Worthington Biochemical Corporation, Cat. LS004188) and 0.25% trypsin (Life Technologies Corporation, Cat. 15400-054) in 1 mL phosphate buffer saline (PBS) at 28℃ for 45 min. After dissociation, supplementation of fetal bovine serum (FBS, Life Technologies Corporation, Cat. 10437028) and centrifugation (600g, 4 min), pellet was resuspended by FACS™ Pre-Sort Buffer (BD Biosciences, Cat. 563503) and filtered through cell strainers. 1 μL Propidium iodide (1 mg/μL) was added to the eluate to stain dead cells. GFP^+^ cells were isolated by FACSAria IIIu–15 color cell sorter (BD Biosciences) and collected into 12.5 μL lysis buffer (Takara Bio USA, Cat. 634439). The cell numbers ranged from 700-1000 cells in *foxl2l*^-/-^ or 300-700 cells in *foxl2l*^+/+^, but only 300 cells were used for further experiments. Then the cell lysates were used for reverse transcription and 10-cycle amplification by SMART-Seq HT kit seq (Takara Bio USA, Cat. 634437) following the user manual. The cDNA generated from cell lysates were quantified by Bioanalyzer (Agilent Technology) to check cDNA concentration and quality. The libraries were generated by Nextera XT (Illumina) and sequenced by NextSeq 500 (Illumina) following the manufacture’s protocols.

The RNAseq fastq data were imported into CLC Genomics Workbench v.10.1.1 (Qiagen) for data trimming, mapping sequence, and differential expression analysis. Raw sequencing reads were trimmed by removing adapter sequences, low-quality sequences (Phred quality score < 20), and sequences with lengths > 30 bp. Sequencing reads were mapped to the zebrafish genome from Ensembl (GRCz11), with the following parameters: length fraction = 0.9, and similarity fraction = 0.9. Gene expressions were based on transcripts per million (TPM). Differential gene expression between wild types and mutants were estimated by generalized linear model [24]. False discovered rate (FDR) adjusted p-value (q-value) < 0.05 and fold change > 2 were used as the criteria of differential expressed genes (DEGs). Gene ontology (GO) enrichment analysis was conducted by Metascape [25]. Down-and up-regulated DEGs were analyzed separately and the *p* < 0.05 was standard to identify enriched GO term. Pathway enrichment was analyzed by DAVID 6.8 [26] using Kyoto Encyclopedia of Genes and Genomes (KEGG) pathway database [27]. The significance of enriched pathway was p < 0.05. The dot-plots of enriched GO terms and KEGG pathways were drawn by ggplot2 [28] in R 4.0.2.

### Quantitation of RNA amounts by RT-qPCR

The trunks of 21-dpf wildtype or *foxl2l*^-/-^ zebrafish were used for RT-qPCR validation. The total RNAs were extracted by Quick-RNA™ Microprep Kit (Zymo Research, Cat. R1050) following the manufacturer’s instructions. The reverse transcription was conducted by the high-capacity cDNA reverse transcriptase kit (Applied Biosystems, Cat. 4368814). RT-qPCR assays for gene expression analysis were quantified by using StepOne Real-Time PCR System (Applied Biosystems) with Power SYBR® Green PCR Master Mix (Applied Biosystems, Cat. 4367659). The relative quantification method (△△Ct method) was used for gene expression quantification. To obtain △Ct, we subtracted the Ct of test gene by the Ct of the house keeping gene *elongation factor 1-alpha* (*ef1α*) in each sample. Then, the △Ct of test gene in each sample was subtracted by the mean △Ct in wild type samples to get △△Ct. The relative expression level of the test gene was shown as 2^-△△Ct. The primer used in qPCR was listed in the primer list (supplementary Table 1). The statistical analysis was conducted by Student’s *t*-test. Differences with p < 0.05 were considered statistically significant.

### Cell culture transfection

COS-1 cells were cultured in Minimum essential Medium (Gibco) supplemented with 10% fetal bovine serum (Gibco). For IF staining, COS-1 cells were cultured on glass slides in 12-well plates and transfected with 500 ng pcDNA3-*foxl2l*-FLAG. After incubation for 24 h, slides were picked up and incubated with 4% PFA to fix cell and followed by IF method described previously. Anti-FLAG antibody (1:1000) (Sigma, Cat. F1804) and anti-mouse-Alexa555 (1:200) (Invitrogen, Cat. 21424) were used as primary and secondary antibody, respectively. For Western blot, transfected cells in 12-well plates were digested by 100 μL Passive Lysis buffer (Promega, Cat. E1941) per well for 15 min. Cell lysates were heated with sample buffer (10% SDS, 250 mM Tris Base, 0.02% Bromophenol Blue, 5% Beta-mercaptoethanol and 30% Glycerol) at 95℃ for 5 min. Protein samples were separated by 10% SDS-PAGE gel under 150V for 70 min and transferred on a PVDF membrane (Millipore) under 200 A for 90 min. Transferred membranes were incubated with Anti-FLAG antibody (1:1000) overnight at 4℃ after 1 hour blocking in 5% milk. The membrane was then washed by PBST three times and incubated with Anti-mouse-HRP (1:2000) for 1 h. Pierce™ ECL Western Blotting Substrate (ThermoFisher Scientific, Cat. 32106) was used for generating chemiluminescent signals.

## Results

### *foxl2l* is expressed in the juvenile gonads and ovaries

We examined the expression of *foxl2l* in gonad development at the juvenile stage in the TL strain zebrafish by whole mount *in situ* hybridization. No signal was detected at 12 hours postfertilization (hpf), 30 hpf and 3 dpf (Fig. 1Ab-d). The *foxl2l* transcript was detected in gonads starting from 8 dpf (Fig.1Ae, 1Af). The expression was stronger in bigger gonads and weaker in smaller gonads at 8 to 12 dpf (Fig. 1Ae-j). Quantitation of signal intensities indicated that the *foxl2l* levels were gradually elevated from 8 to 18 dpf (Fig. 1B). Among 11 different adult tissues *foxl2l* transcripts were detected by RT-PCR only in the ovary (Fig. 1C). This result indicates that *foxl2l* is a female-specific transcript at the adult stage.

**Fig. 1.**
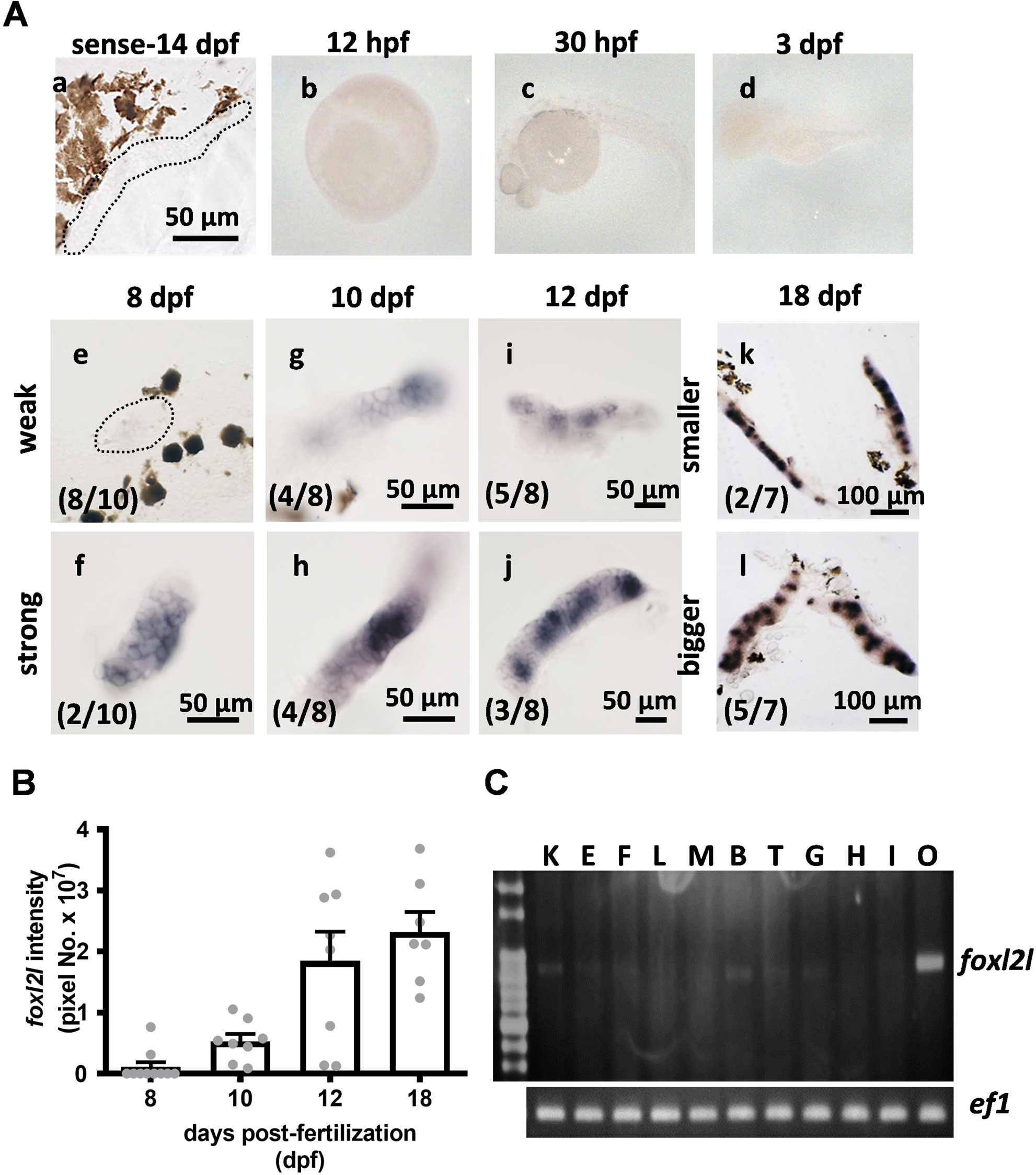
Temporal and sex-dimorphic expression of zebrafish *foxl2l* in the ovary. **(A)**. Whole mount *in situ* hybridization of fish (TL strain) showing *foxl2l* transcripts start at 8 dpf in some gonads. The numbers inside the brackets are the number of gonads showing the staining pattern out of the total number of gonad. The dotted lines outline gonads that are not stained while neighboring pigments are dark brown. **(B).** Quantitation of *foxl2l* intensity. **(C).** RT-PCR analysis of tissue distribution of *foxl2l* transcripts. B, brain; E, eye; F, fin; G, gill; H, heart; I, intestine; K, head kidney; L, liver, M, muscle; O, ovary; T, testis.

### *foxl2l* is expressed in proliferating progenitor germ cells

We further examined the cell type in zebrafish gonads that expressed *foxl2l* by double staining with different cellular markers. At 12 dpf, *foxl2l*^+^ was expressed in some *vasa*^+^ germ cells indicating that *foxl2l* is expressed in a subset of germ cells (Fig. 2Aa). When co-stained with *nanos2*, a germline stem cell marker, *nanos2^+^* cells and *foxl2l^+^* cells were distinct, indicating that *foxl2l-*expressing cells are not germline stem cells (Fig. 2Ab).

Germ cells divide to form cysts, and their DNAs duplicate in cystic cells. All *foxl2l^+^* cells at 25 dpf contained EdU, which could be incorporated into replicating DNA of proliferating cells (Fig. 2Ac). In contrast, some EdU^+^ germ cells did not express *foxl2l*. Moreover, no *foxl2l* existed in EdU^-^ germ cells. These results imply that *foxl2l* is present in proliferating germ cells inside cysts.

### *foxl2l* disappears after germ cells enter meiosis

To observe whether zebrafish female meiotic germ cells express *foxl2l*, the morphology of germ cells were examined. In adult ovary, *foxl2l* appeared in oogonia that have a single nucleolus in the center of the nucleus (Fig. 2Ba). However, *foxl2l* disappeared at zygotene (nucleolus at periphery), pachytene (individual cell with a peripheral nucleolus and chromatin condensed with weak DAPI staining) and diplotene (weak DAPI staining and surrounded by somatic cells) stage of germ cells (Fig. 2Bb-d). Therefore, *foxl2l* is expressed in pre-meiotic germ cells and disappeared when germ cell enters zygotene stage of prophase meiosis I.

### Foxl2l is located in the nucleus

To further identify the subcellular locations of Foxl2l, plasmid (pcDNA3-*foxl2l*-FLAG) overexpressing Foxl2l with FLAG tag was transfected into COS-1 cells (Fig. 2C). Immunoblot analysis indicated that Foxl2l-FLAG (31 kDa) was detected in the cells transfected with pcDNA3-*foxl2l*-FLAG but not with pcDNA3-FLAG (Fig. 2D). Immunofluorescent image showed that Foxl2l-FLAG was co-located with nuclear DAPI staining (Fig. 2E). This result indicates that Foxl2l is a nuclear protein.

**Fig. 2.**
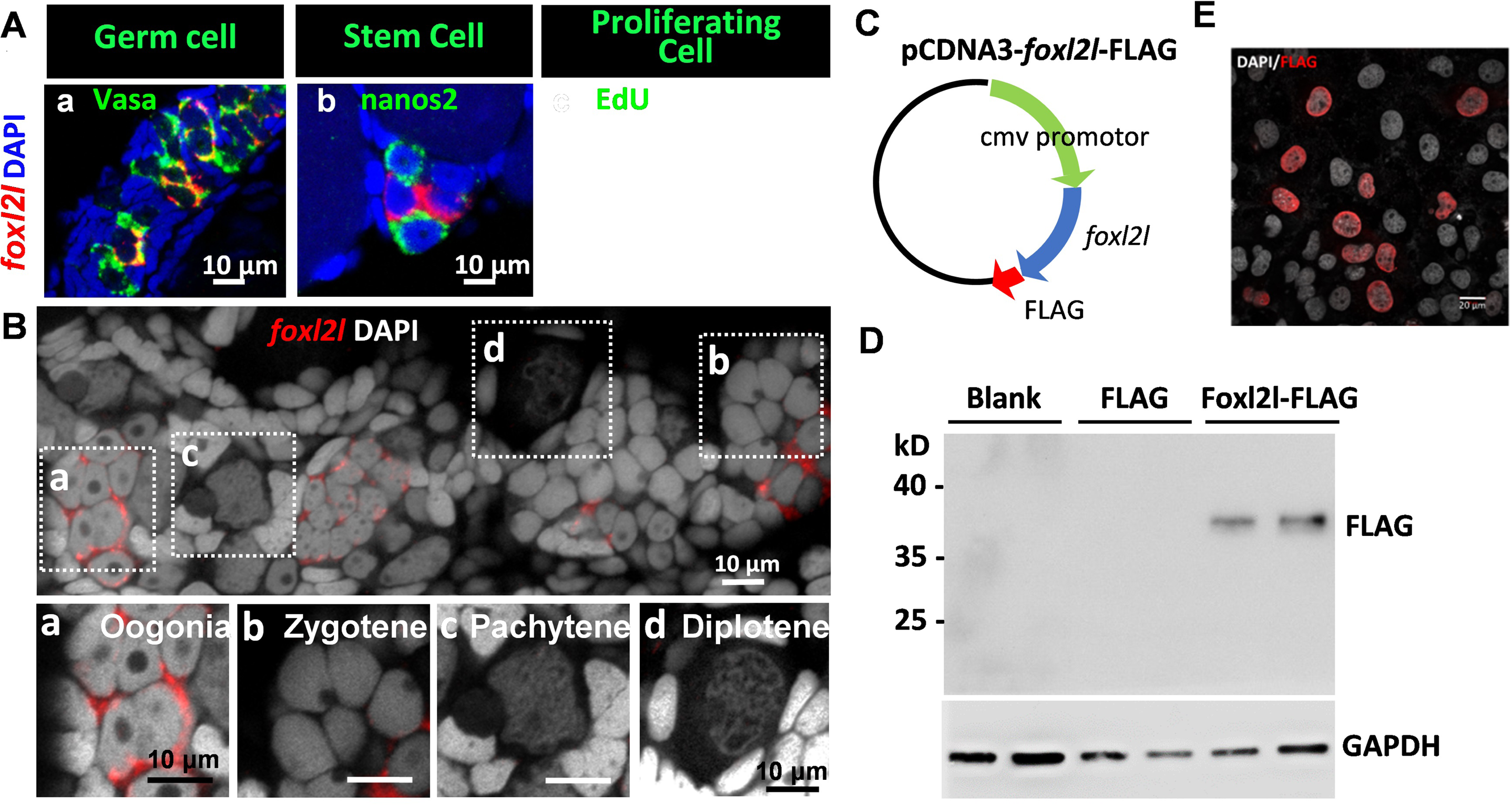
Foxl2l is a nuclear protein, and its transcript presents in proliferating germ cells disappears at meiosis. **(A)**. The *foxl2l* transcript detected by whole mount fluorescence *in situ* hybridization (red) is co-stained with markers for (a) germ cell (*vasa*) and (b) germline stem cell (*nanos2*) showing it is present only in non-stem germ cells. (c) The *foxl2l* is present in a subset of EdU^+^ cells (labeled by an asterisk), and absent in other EdU^+^ cells (arrow). **(B)**. Lack of *foxl2l* expression in germ cells at the meiotic prophase. The *foxl2l* signals are detected in a subset of oogonia, but not in meiotic cells (zygotene, pachytene, diplotene) in adult ovary. **(C)**. The map of pCDNA3-*foxl2l*-FLAG plasmid. **(D)**. Immunoblot proteins in COS-1 cells overexpressing nothing (Blank), FLAG or *foxl2l*-FLAG plasmid using FLAG antibody. Every sample is loaded in two lanes. **(E).** Immunofluorescence detects FLAG antibody staining in the nucleus of COS-1 cells transfected with pCDNA3-*foxl2l*-FLAG plasmid.

### Mutation of *foxl2l* leads to testis formation

In zebrafish, *foxl2l* gene only contains one exon that encodes a forkhead (FH) domain. Three gRNAs were designed in three different target sites (1) 5’-untranslated region, (2) N-terminal region before the FH domain, and (3) the beginning of FH domain. These gRNAs were injected into embryos to generate F0 founders that later breed to form *foxl2l* deficient lines (Fig. 3A). Normal adult females possess genital papilla that is absent in the males (Fig. 3B). All of the *foxl2l^-/-^* fish, however, lack genital papilla. These fish all contain testes instead of ovaries. Histological staining also showed that *foxl2l^-/-^* gonads possessed testes containing sperm cells at all stages.

**Fig. 3.**
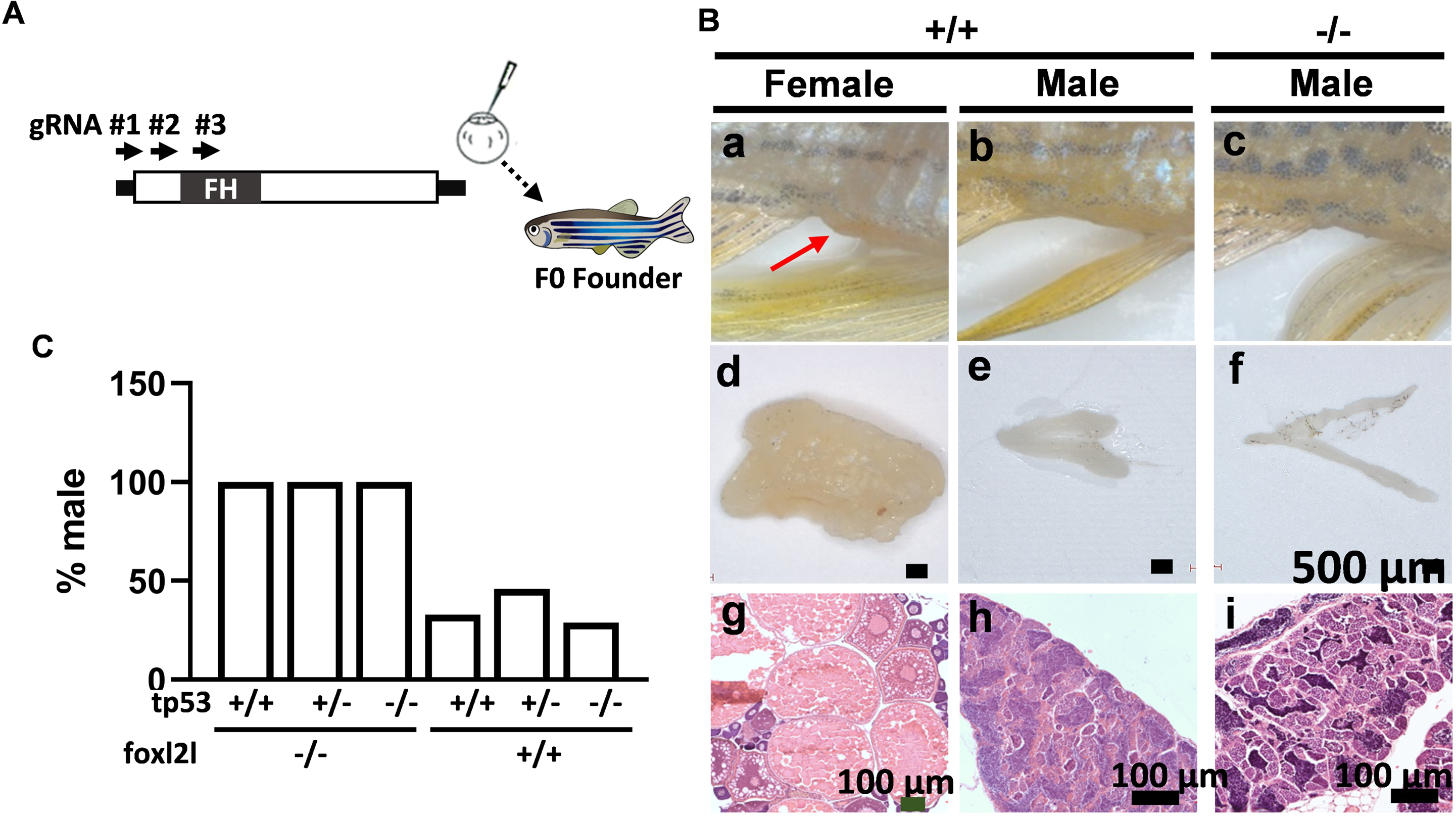
Disruption of *foxl2l* leads to testis formation unrelated to *tp53*. **(A).** The target sites of the gRNAs for *foxl2l* (arrows) and their injection into the F0 embryos. **(B).** The morphology of adult fish. (a-c) the ventral belly, (d-f) dissected gonad, and (g-i) HE staining after sectioning. The red arrow points to the genital papilla present in the female fish. **(C).** The ratios of male fish in *tp53* and *foxl2l* single and double mutant fish. The *foxl2l*^-/-^*tp53^-/-^* mutants have the same sex ratios as *foxl2l^-/-^tp53^+/+^* fish showing that *tp53* does not influence sex ratios.

Apoptosis of oocytes is a critical process to switch female fate to male fate in zebrafish during juvenile stage [5]. In some zebrafish mutants, all-male phenotypes are mediated by oocyte apoptosis and the sex ratio can be reverted by depleting *tp53* [8, 29, 30]. To examine whether the induced masculinization in *foxl2l* mutants is mediated by apoptosis, *foxl2l* mutant fish were mated with *tp53* mutant fish to generate double mutant fish. The all-male phenotype of *foxl2l* mutants was not altered when coupled with *tp53* heterozygous or homozygous mutations (Fig. 3C). This result shows that the masculinization process in *foxl2l* mutant is unrelated to oocyte apoptosis.

### *foxl2l*:EGFP fish mark *foxl2l^+^* cells by GFP

To investigate the expression of *foxl2l* during gonad development in more detail, we used the tol2 system to generate transgenic fish that express EGFP in the *foxl2l^+^* cells. The *foxl2l* proximal promoter fragment from -3048 to +14 of *foxl2l* was isolated from genome by PCR and cloned into pT2KXIG plasmid for the generation of transgenic fish (Fig. 4A). At 16 dpf, the EGFP was well co-localized with *foxl2l* mRNA (Fig. 4A). It indicates that a *foxl2l* promoter (−3048 to +14) is sufficient to drive the expression of EGFP in the gonad of transgenic fish.

**Fig. 4.**
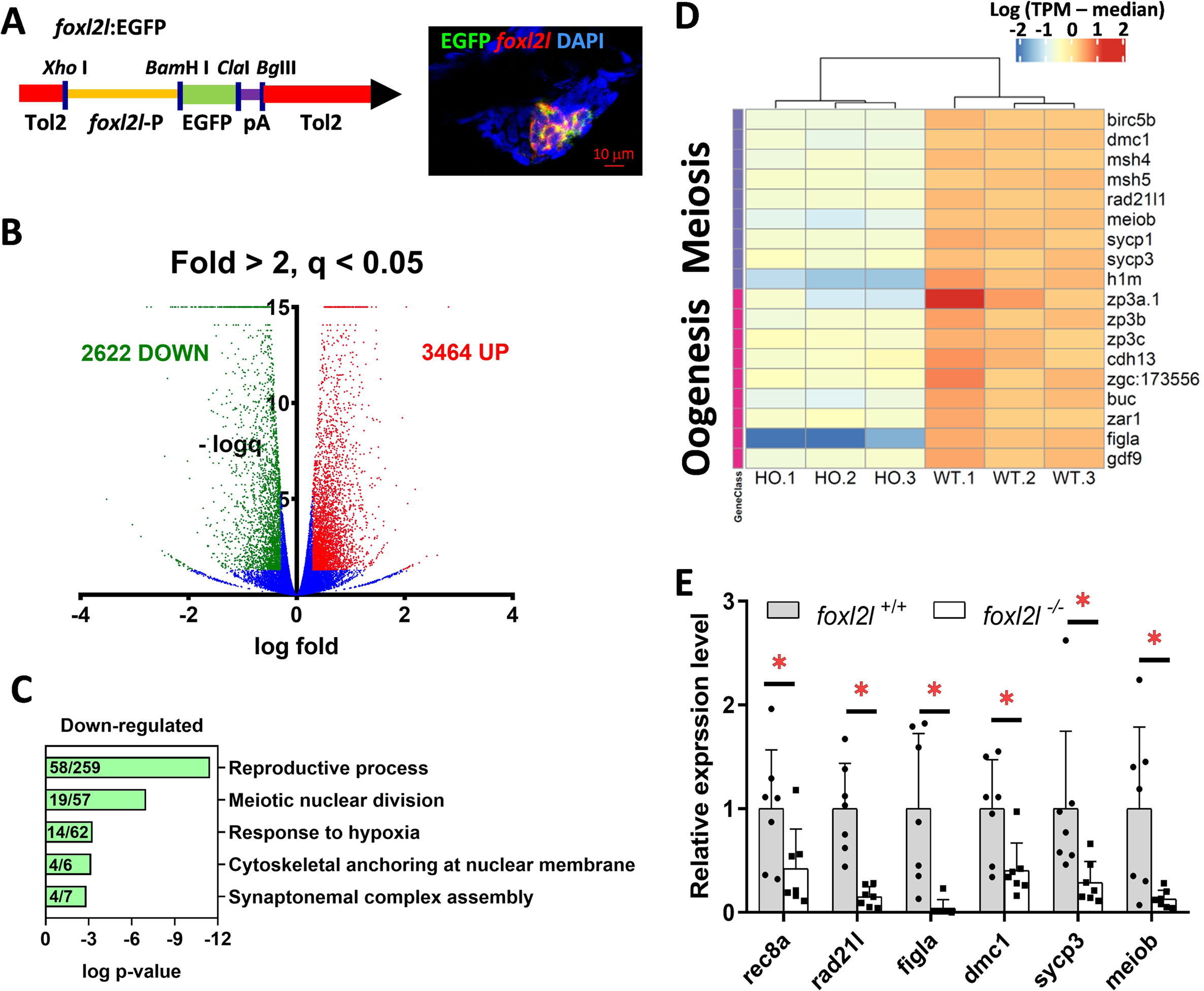
RNA-seq reveals downregulation of genes in meiosis and oogenesis in *foxl2l* mutants. **(A)**. Map of the plasmid used for the generation of transgenic fish expressing EGFP in cells expressing *foxl2l*. A fragment of *foxl2l*-P promoter (−3048 to +14) drives the expression of EGFP linked to poly A (pA). Double staining (right figure) shows EGFP (green) and *foxl2l* (red) are expressed in the same cell. **(B).** Volcano plots of differentially expressed genes (DEGs). DEG is defined as fold change>2 and false discovery rate adjusted p value (q-value)<0.05. Total 3464 and 2622 genes were upregulated (red) and downregulated (green) in *foxl2l^-/-^* cells, respectively. **(C).** Enrichment analysis of gene ontology among downregulated genes in *foxl2l* mutant cells. The numbers inside the bar indicate the ratio of DEGs to the number of all genes in this GO term. Reproductive process (GO:0022414), meiotic nuclear division (GO:0140013), response to hypoxia (GO:0001666), cytoskeletal anchoring at nuclear membrane (GO:0090286), and synaptonemal complex assembly (GO:0007130). **(D).** Heatmap of genes in meiosis and oogenesis from three batches of wildtype (WT) and *foxl2l* homozygous mutant (HO) cells. **(E).** qPCR detects down-regulation of meiotic genes in 21-dpf *foxl2l^-/-^* fish.

### Downregulation of genes in meiosis and oogenesis in *foxl2l* mutant

To examine the changes of gene expression caused by *foxl2l* deficiency, EGFP-expressing germ cells were collected from *foxl2l*:EGFP transgenic fish on *foxl2l*^-/-^ or *foxl2l*^+/+^ background at 21 dpf for bulk RNA-seq analyses (supplementary Fig. S1). The principal components of WT and mutant groups were well separated (supplementary Fig. S2), indicating the reliability of the analysis. With the criteria of fold change > 2 and q-value < 0.05, we identified 2,622 down-regulated and 3,464 up-regulated genes in *foxl2l* mutants (Fig. 4B). Among down-regulated genes, the most enriched GO terms were reproductive process and meiotic nuclear division (Fig. 4C). We chose many genes in meiosis and oogenesis for heatmap analysis and found that they were downregulated in *foxl2l* mutant (Fig. 4D). qPCR analysis also showed that meiotic genes (*rec8a*, *sycp3*, *rad21l*, *dmc1* and *meiob*) and oogenic gene (*figla*) were repressed in the *foxl2l* mutant (Fig. 4E).

### *Foxl2l*:EGFP+ cells were accumulated in *foxl2l* mutant

We examined *foxl2l*-expressing cells by immunostaining (Fig. 5). At 21 dpf, *foxl2l*:EGFP was expressed exclusively in proliferating cystic cells and absent in GSCs and meiotic germ cells (Fig. 5Aa-c), same as the expression of *foxl2l* mRNA. This further confirms the usefulness of this transgenic line in tracking *foxl2l*-expressing cells.

**Fig. 5.**
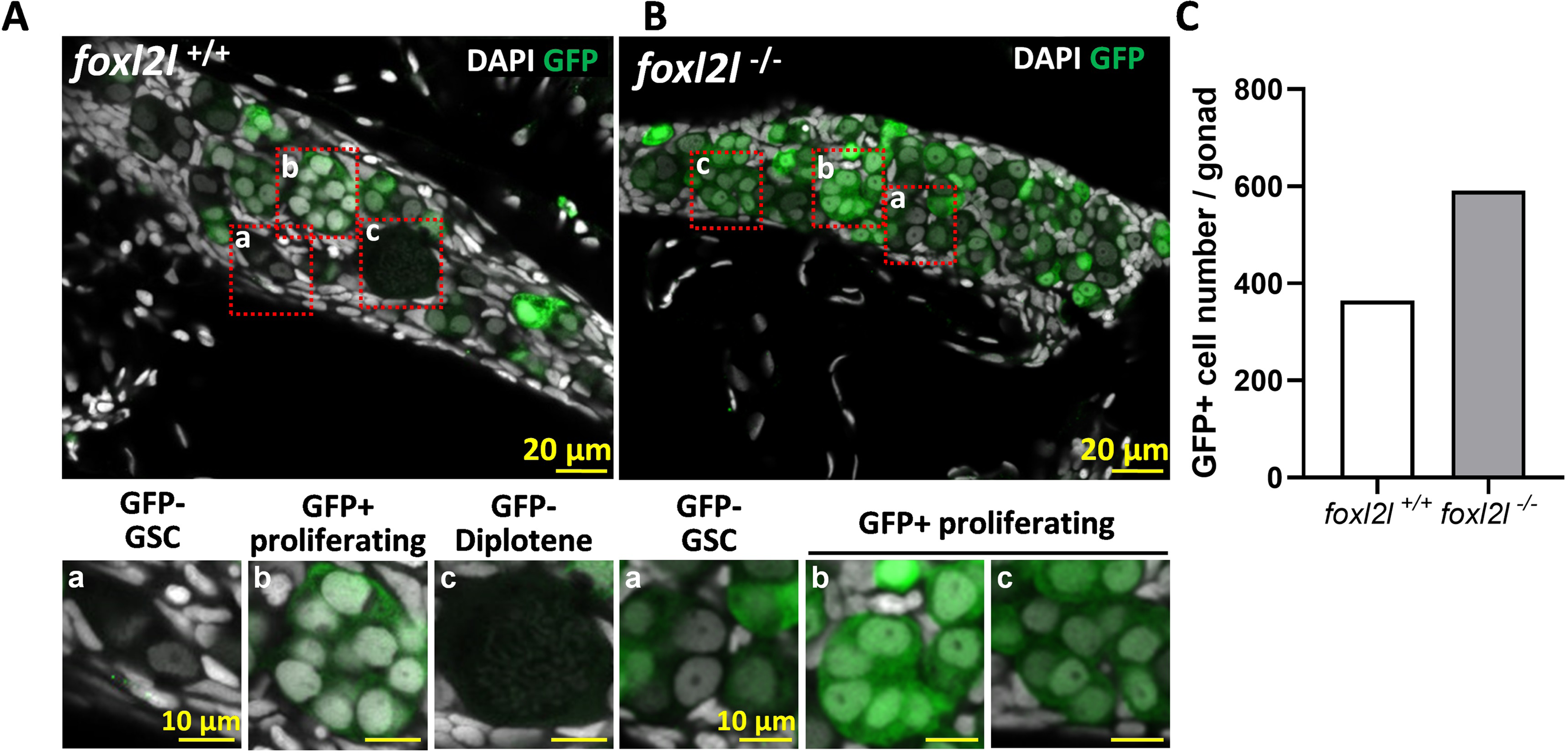
Accumulation of germ cells after *foxl2l* depletion. **(A-B)**. Staining of 21-dpf gonads with EGFP and DAPI. The developmental stages of germ cells were recognized by different patterns of DAPI staining. Germline stem cells are small and EGFP^-^. Proliferating cells are small, EGFP^+^, and located in cysts. Diplotene germ cells are large and DAPI^-^. EGFP is present only in cysts, but not in GSC and diplotene oocytes of WT fish. The *foxl2l^-/-^* gonads lack meiotic oocytes and accumulate EGFP^+^ progenitors. **(C).** Cell counting indicates that EGFP^+^ cells are accumulated in mutant gonads.

In *foxl2l^-/-^* fish, many EGFP^-^ GSCs and EGFP^+^ cystic proliferating cells were observed, but meiotic germ cells were missing (Fig. 5B). This result indicates that *foxl2l* is required to meiotic entry. We counted the number of EGFP^+^ cells and found that more EGFP^+^ cells were present in *foxl2l*^-/-^ than WT fish at 21 dpf (Fig. 5C). This result indicates the accumulation of *foxl2l*-expressing cells in the *foxl2l* mutant.

### Novel pathways were altered in *foxl2l* mutant

Our pathway enrichment analysis shows that in addition to meiosis and oogenesis, other pathways are also altered in the *foxl2l* mutants (supplementary Fig. S3). The genes upregulated in *foxl2l* mutants are enriched in pathways for primary bile acid biosynthesis, mismatch repair, and biosynthesis of unsaturated fatty acids (supplementary Fig. S4). Conversely, genes down-regulated in *foxl2l* mutants are enriched in pathways for phagosome, spliceosome, and Wnt signaling (supplementary Fig. S5).

We tested the expression of individual genes in these pathways both by RNAseq and by q-PCR. Genes for bile acid synthesis such as *cholesterol 25-hydroxylase-like protein 1 member 1* (*ch25hl1.1*) and *acyl-coenzyme A thioesterase 8* (*acot8*), in DNA mismatch repair such as *mutL homolog 1* (*mlh1*) and *replication protein A1* (*rpa1*), and in the synthesis of unsaturated fatty acids such as *elongation of very long chain fatty acids protein 2* (*elovl2*) and *fatty acid desaturase 2* (*fads2*) were all up-regulated in the *foxl2l* mutant (Fig. 6). On the contrary, the gene for phagosome and spliceosome, *ATPase H^+^ transporting V0 subunit ca* (*atp6v0ca*) and *small nuclear ribonucleoprotein polypeptide G* (*snrpg*), respectively, two Wnt signaling genes, *wnt8a* and *wnt5b*, were down-regulated in the *foxl2l* mutant (Fig. 6).

**Fig. 6.**
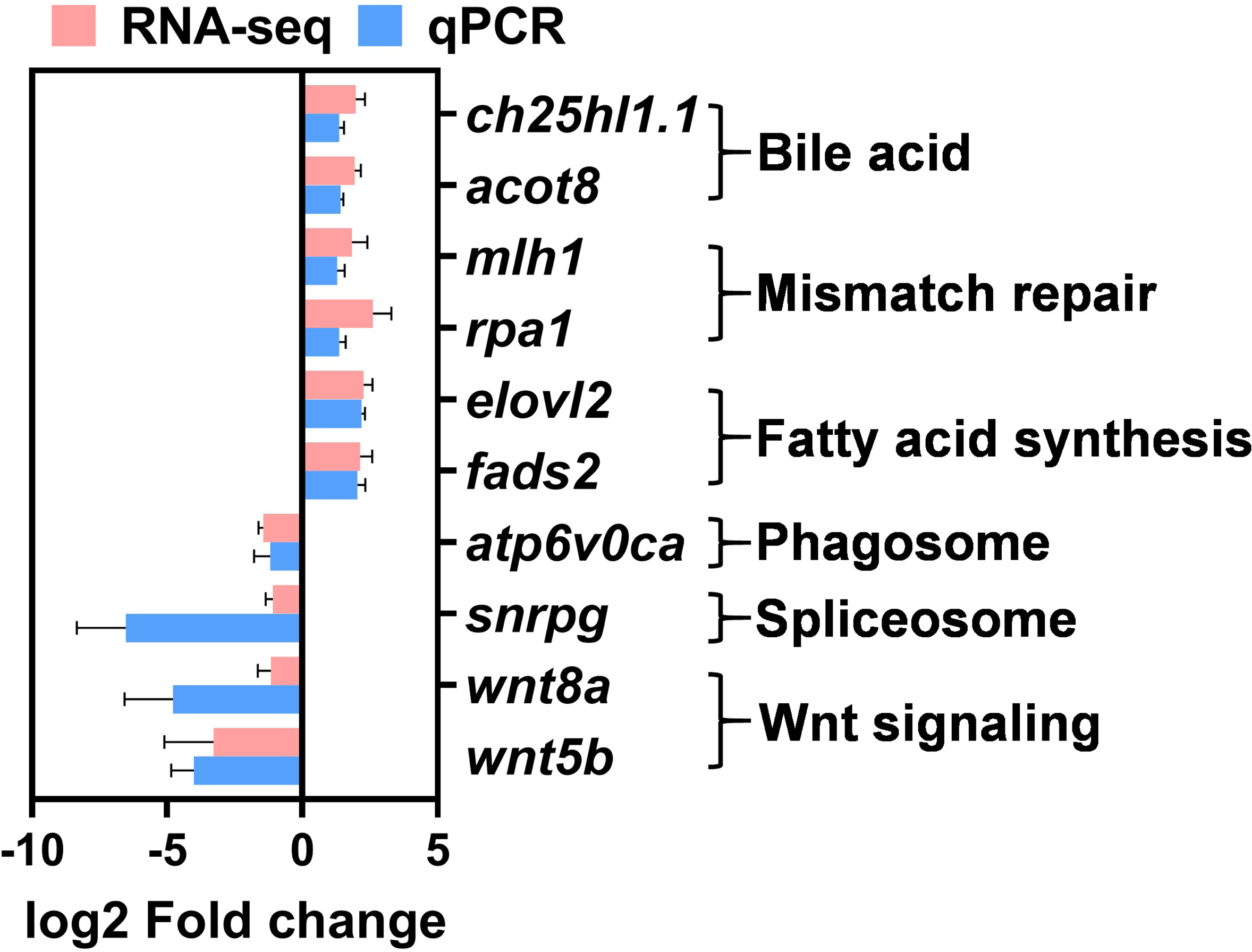
DEGs were enriched in novel pathways. The Log-2-fold changes of DEGs in enriched pathways analyzed by RNA-seq and qPCR.

## Discussion

### Comparison of *foxl2l* in different species

Foxl2l has also been identified in other teleost species. In teleost including rainbow trout (*Oncorhynchus mykiss*), killifish (*Kryptolebias marmoratus*) and Nile tIilapia (*Oreochromis niloticus*), *foxl2l* is highly enriched in female gonad than male gonad, which resemble the observation in zebrafish and medaka [31–33]. However, *foxl2l* is predominant in male gonad in some other teleosts, including protogynous fish, ricefield eel (*Monopterus albus*) and orange-spotted grouper (*Epinephelus coioides*), and gonochoristic fish, Atlantic salmon (*Salmo salar*), Japanese eel (*Anguilla japonica*) and European sea bass (*Dicentrarchus labrax*), suggesting the importance of Foxl2l in male gonad differentiation [34–38]. The differential expression among different species reflects the divergent roles of Foxl2l, perhaps due to the highly variable C-terminal region among different species [38]. Here we show zebrafish *foxl2l* plays roles important in ovaries rather than testes.

### *foxl2l* is expressed in the proliferating cells but not GSC and oocyte

In medaka, *foxl3* signals are detected in all cystic mitotic germ cells and a subset of self-renewed mitotic germ cells, but not in meiotic germ cells [17]. In zebrafish, we show here that *foxl2l* is expressed in mitotic germ cells differentiation from germline stem cells and before oocytes. Furthermore, *foxl2l* is only detected in EdU^+^ proliferating cells in both species. The types of *foxl2l-*expressing cells in medaka and zebrafish are similar. These expression results are consistent with our function data that *foxl2l* is involved in oogenesis at the onset of meiosis.

### *foxl2l* promotes female fate unrelated to germ cell death

Without foxl2l, zebrafish larvae lack oocytes and develop as males. Degeneration of oocytes at the juvenile stage is a route for male development in zebrafish [5]. The inhibition of oocyte degeneration by mutation of *tp53* can rescue all-male phenotypes in some zebrafish mutants [30]. However, we show here that the all-male phenotype in *foxl2l* mutants cannot be rescued by depleting *tp53*. Different from other mutants, oocytes are absent in the *foxl2l^-/-^* gonads and we fail to detect massive germ cell apoptosis. These results indicate that *foxl2l* mutation leads to a change of germ cell fate unrelated to germ cell apoptosis

### Comparison of zebrafish and medaka *foxl2l* orthologues

Our gene expression analysis and IF images show that *foxl2l^-/-^* larval germ cells do not express meiotic and oogenic genes, and stay at the stage before meiotic entry. These results indicate that *foxl2l* is required for germ cells to express meiotic and oogenic genes for entering oogenesis. In medaka, *foxl3* (*foxl2l* ortholog) directly activates meiotic gene *rec8a* and oogenic gene *fbxo47* and *figla* [18]. The *fbxo47* orthologue is not present in zebrafish, but *fbxo47* downstream gene *figla* and of oogenic genes are downregulated in the zebrafish *foxl2l* mutant. Thus, *foxl2l* plays similar roles in promoting oogenesis in zebrafish and medaka although their action mechanisms may not be identical.

Although zebrafish *foxl2l* and medaka *foxl3* share similar expression pattern and functions, the phenotypes of their mutation are not identical. Zebrafish *foxl2l* mutants are all males with male secondary sex characteristics. Medaka XX *foxl3* mutants, however, are females with female secondary sex characteristics although their ovary-like gonads which contain fertile sperms [17]. The difference between the phenotypes of the same orthologous *foxl2l* gene in zebrafish and medaka may be due to the difference of sex determination in these two species.

Zebrafish laboratory strains lack sex chromosomes [39], and environmental factors such as density, temperature, and food availability play important roles for their sex differentiation [1, 40, 41]. The sex of medaka is basically determined by the sex-determining gene, *DM-domain gene on the Y chromosome* (*dmy*) [42], although environmental factors can alter the sex differentiation processes [43–45]. The XY gonad of medaka contains *dmy*, which promotes male development and suppresses female germ cell proliferation [3]. Medaka XX gonads do not contain *dmy*, and their XX gonads differentiate into ovary even in the absence of *foxl3.* On the contrary, zebrafish *foxl2l^-/-^* mutants fail to develop into females and 100% of them become males.

### Pathways upregulated in *foxl2l* mutants

We detected upregulation of genes in the mismatch repair pathway. Besides DNA repair, components of mismatch repair are also involved in meiosis. The MutL homolog (MLH) complexes play important roles in meiotic recombination [46]. Depletion of *Mlh1* in mice causes infertility and meiotic arrest at the stage of spermatocytes [47]. The genes for homologous recombination, such as *dmc1* and *rad51* were downregulated. However, genes for mismatch repair, *mlh1* and *rpa1*, were up-regulated in the *foxl2l^-/-^* germ cells in our RNAseq and q-PCR data. Our single-cell RNA-seq database reveals that *mlh1* is highly expressed in all progenitor germ cells and *rpa1* is highly expressed in the early progenitors (Prog-E) (NCBI GEO, GSE173718). Whereas *dmc1* and *rad51* are expressed in meiotic oocytes (Clusters 8 to 10) [21]. We show here that progenitor cells are accumulated in the *foxl2l* mutants; therefore it makes sense that progenitor transcripts such as *mlh1* and *rpa1* are upregulated. On the contrary, because mutant cells fail to enter meiosis, meiotic genes such as *dmc1* and *rad51* are absent in the *foxl2l* mutants. Our gene expression data is consistent with the stage of germ cells in the mutants.

We detected increased expression of genes involved in the synthesis of unsaturated fatty acid. This indicates that the amount of lipid may be increased in the *foxl2l^-/-^* germ cells. Control of lipid homeostasis may be important for sex differentiation. Inhibition of fatty acid synthesis leads to female-to-male sex reversal in medaka [45]. The involvement of fatty acid in zebrafish sex differentiation is another subject that needs to be explored.

### Alternative downregulated pathways in *foxl2l* mutant

In addition to meiosis and oogenesis, we detected down regulation of pathways involved in spliceosome and phagosome. The functions of spliceosome and phagosome in germ cells have been rarely studied. Aberrant spliceosome expression and altered alternative splicing events correlate with maturation deficiency in human oocytes [48].

We also detected downregulation of pathways in Wnt signaling in *foxl2l* mutants. Wnt signaling is important for ovary differentiation [49]. *Wnt5b* has higher expression levels in ovaries over testes in Chinese soft-shelled turtle (*Pelodiscus sinensis*), Muscovy duck (*Cairina moschata*) and Catfish (*Clarias batrachus*) [50–52], therefore *Wnt5b* may participate in the differentiation of ovaries. In zebrafish we show here that *wnt5b* is down-regulated in *foxl2l* mutants; thus zebrafish *wnt5b* may be a female gene involved in ovary development.

## Conclusion

Here we have characterized a transcription factor, Foxl2l, which plays key roles in ovary differentiation. Foxl2l is present exclusively in the mitotic germ cells. The depletion of *foxl2l* leads to the accumulation of mitotic germ cells and a block to enter meiotic oogenesis. We have characterized expression of genes during this process that finally leads to female-to-male sex reversal in the mutants. We show here that *foxl2l* is a gene in the mitotic germ cells important for female fate.

## Supplementary Data

### Funding

This work was funded by grants from Academia Sinica, AS-101-TP-B05, NHRI-EX107-10506SI, MOST 107-2321-B-001-034, MOST 108-2311-B-001-038-MY3.

### Acknowledgments

We would like to thank Taiwan Zebrafish Core Facility, Genomics Core, Bioinformatics Core and Imaging Core (Institute of Molecular Biology Academia Sinica) for help in confocal imaging and transcriptome data analysis. Academia Sinica Core Facility and Innovative Instrument Project (AS-CFII108-113) for cell sorting service.

### Author contribution

CHY performed mutant analysis and RNAseq experiments, and analyzed the data. YwW examined *foxl2l* expression and generated *foxl2l* mutant fish lines. CwH performed the *tp53* experiment, and YJP initiated the germ cell proliferation and germ cell counting experiments. BcC secured funding, designed the experiments, and oversaw the execution of the project. CHY, CwH, and BcC wrote the initial draft. Everyone participated in the revision of the manuscript.

## Supplementary Information

**Supplementary Table S1.**
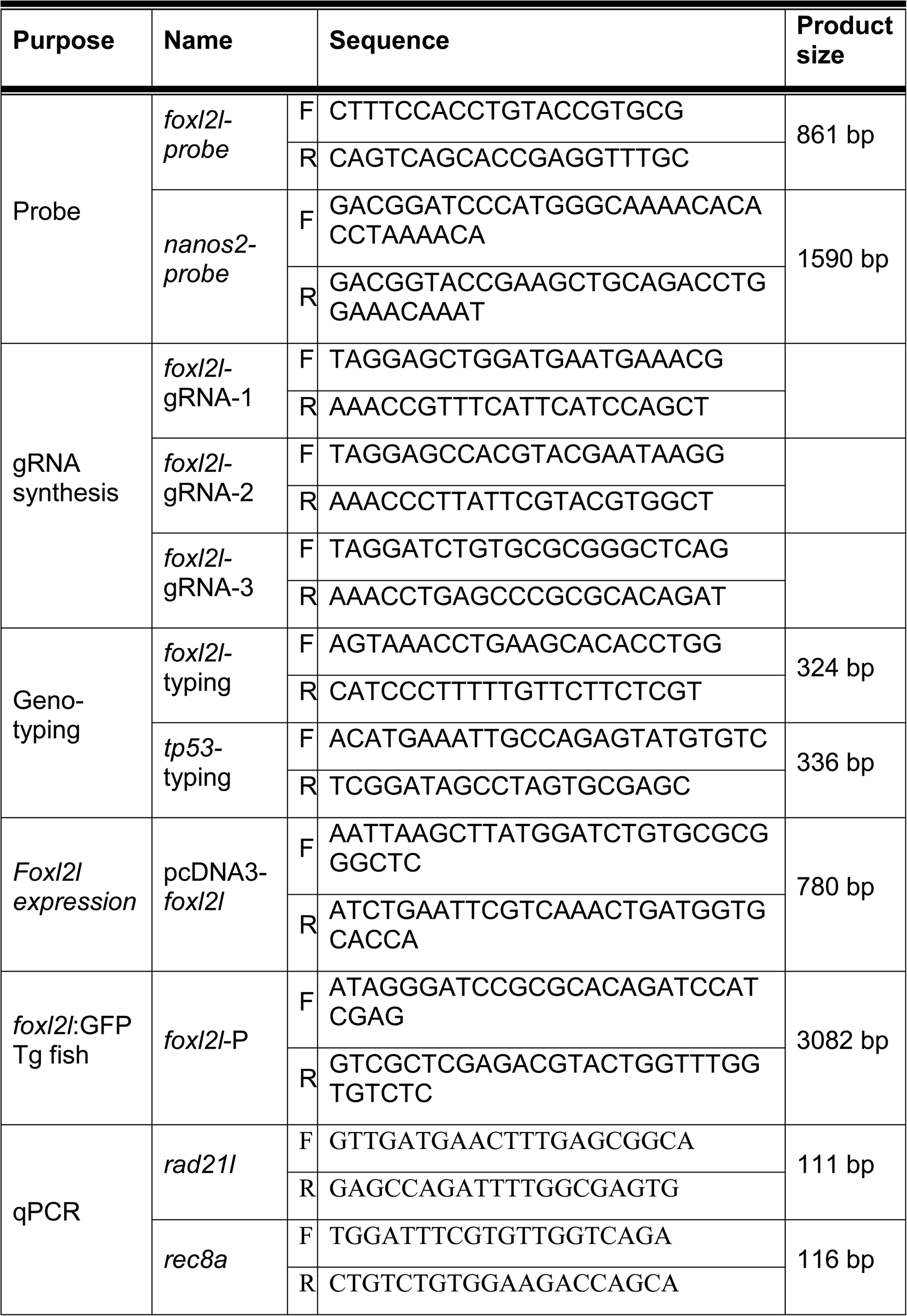
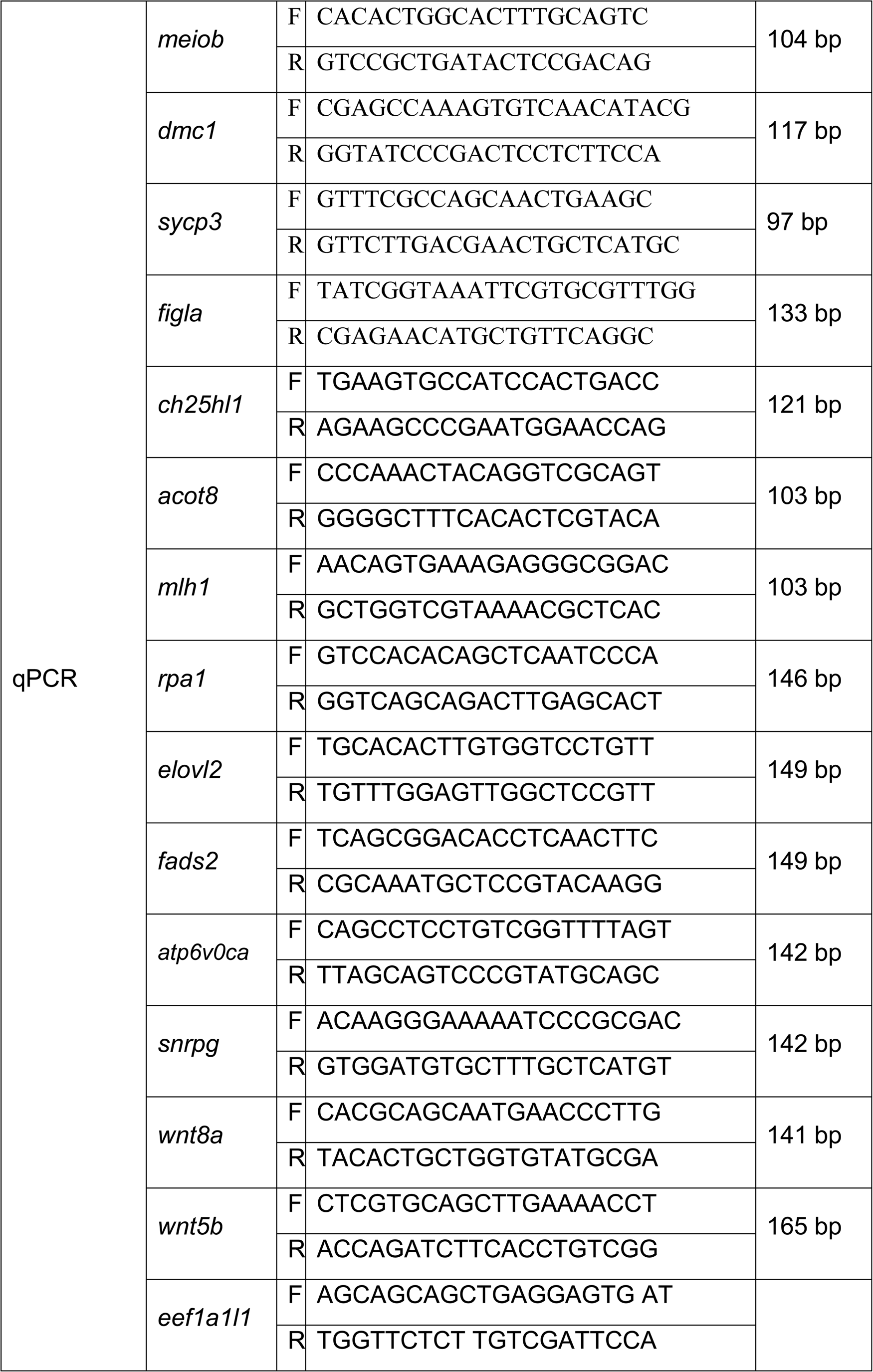
Sequences of oligonucleotide primers used in this work.

**Supplementary Table S2.** The list of plasmids used in this work.

| Purpose | Plasmid | Insert description | Source |
| --- | --- | --- | --- |
| *foxl2l* probe | pGEM-Teasy-*foxl2l* | *foxl2l*<br>-56 to +805 | This study |
| *vasa* probe | pTY27-*vasa* | *vasa*<br>+1728 to +2865 | {Yoon, 1997 #1397} |
| *nanos2* probe | pCS2<sup>+</sup>-*nanos2* | *nanos2*<br>-34 to 1556 | This study |
| *transposase* exprssion | pCS-TP | *transposase*<br>Full length | {Kawakami, 2004 #811; Kawakami, 2004 #1299} |
| *foxl2l* gRNA #1 | pT7-gRNA- *foxl2l* #1 | *foxl2l*<br>-20 to -1 | This study |
| *foxl2l* gRNA #2 | pT7-gRNA- *foxl2l* #2 | *foxl2l*<br>+3 to +22 | This study |
| *foxl2l* gRNA #3 | pT7-gRNA- *foxl2l* #3 | *foxl2l*<br>+109 to +128 | This study |
| *foxl2l*:GFP Tg fish | pT2KXIG- *foxl2l*:EGFP | *foxl2l*<br>-3048 to +14 | This study |
| *foxl2l* expression | pcDNA3-*foxl2l*-FLAG | *foxl2l*<br>+1 to +780 | This study |

## Supplementary Figure Legend

**Fig. S1. Collecting *foxl2l*:EGFP^+^ cells through FACS.** FACS plots of sorting *foxl2l*:EGFP+ cells from **(A-E).** wild type or **(F-J).** *foxl2l* KO fish with *foxl2l*:EGFP transgene.

**Fig. S2. The basic analysis of RNA-seq samples. (A).** Mapping rate of reads in each sample. **(B).** Principle analysis of six independent samples showing that WT and HO samples are well separated. **(C).** Cluster analysis of six independent samples showing that WT and HO samples cluster with themselves but not with each other. **(D).** MA plot of DEGs.

**Fig. S3. Pathway enrichment analysis of DEGs between wildtype and *foxl2l* mutant germ cells.** The numbers inside the bar indicate the ratio of DEGs to the number of all genes in this pathway.

**Fig. S4. Up-regulated DEG enriched pathways. (A).** Primary bile acid biosynthesis (dre00120). **(B).** Mismatch repair (dre03430) **(C).** Biosynthesis of unsaturated fatty acids (dre01040). DEGs in the pathways are indicated by red stars.

**Fig. S5. Down-regulated DEG enriched pathways. (A).** Phagosome (dre04145). **(B).** Spliceosome (dre03040). **(C).** Wnt signaling pathway (dre04310). DEGs in the pathways are indicated by red stars.

