## supplementary figures and legends for "Zebrafish Foxl2l functions in proliferating germ cells for female meiotic entry"

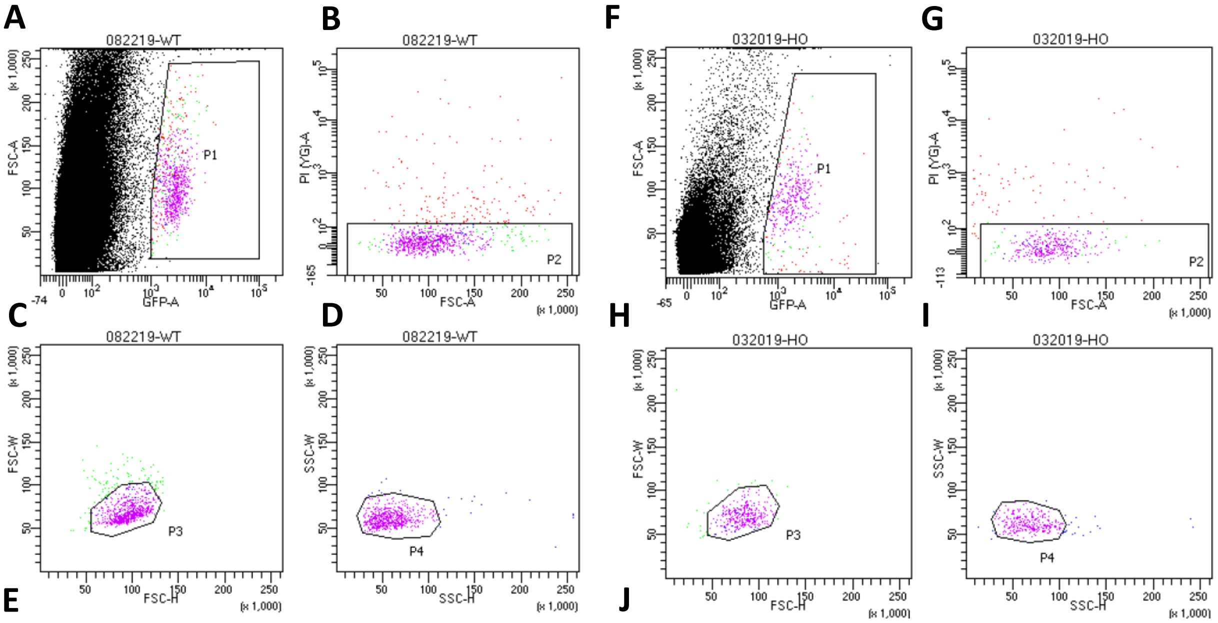


**E**

Tube: WT

| Population | #Events | %Parent | %Total |
|---|---|---|---|
| All Events | 756,084 | #### | 100.0 |
| P1 | 925 | 0.1 | 0.1 |
| P2 | 770 | 83.2 | 0.1 |
| P3 | 674 | 87.5 | 0.1 |
| P4 | 649 | 96.3 | 0.1 |
| P5 | 407,546 | 53.9 | 53.9 |

**J**

Tube: HO

| Population | #Events | %Parent | %Total |
|---|---|---|---|
| All Events | 69,889 | #### | 100.0 |
| P1 | 417 | 0.6 | 0.6 |
| P2 | 348 | 83.5 | 0.5 |
| P3 | 324 | 93.1 | 0.5 |
| P4 | 295 | 91.0 | 0.4 |
| P5 | 41,600 | 59.5 | 59.5 |

**Fig. S1. Collecting *foxl2l*:EGFP⁺ cells through FACS.** FACS plots of sorting *foxl2l*:EGFP+ cells from **(A-E).** wild type or **(F-J).** *foxl2l* KO fish with *foxl2l*:EGFP transgene.

**A**

| | WT-1 | WT-2 | WT-3 | HO-1 | HO-2 | HO-3 |
|---|---|---|---|---|---|---|
| Total reads | 101,620,650 | 116,154,326 | 96,354,520 | 104,751,262 | 108,488,956 | 93,468,992 |
| Total mapped reads | 87% | 83% | 64% | 87% | 86% | 88% |
| Unique mapped reads | 84% | 80% | 62% | 84% | 83% | 86% |
| Read mapped Gene | 77% | 72% | 56% | 76% | 74% | 80% |

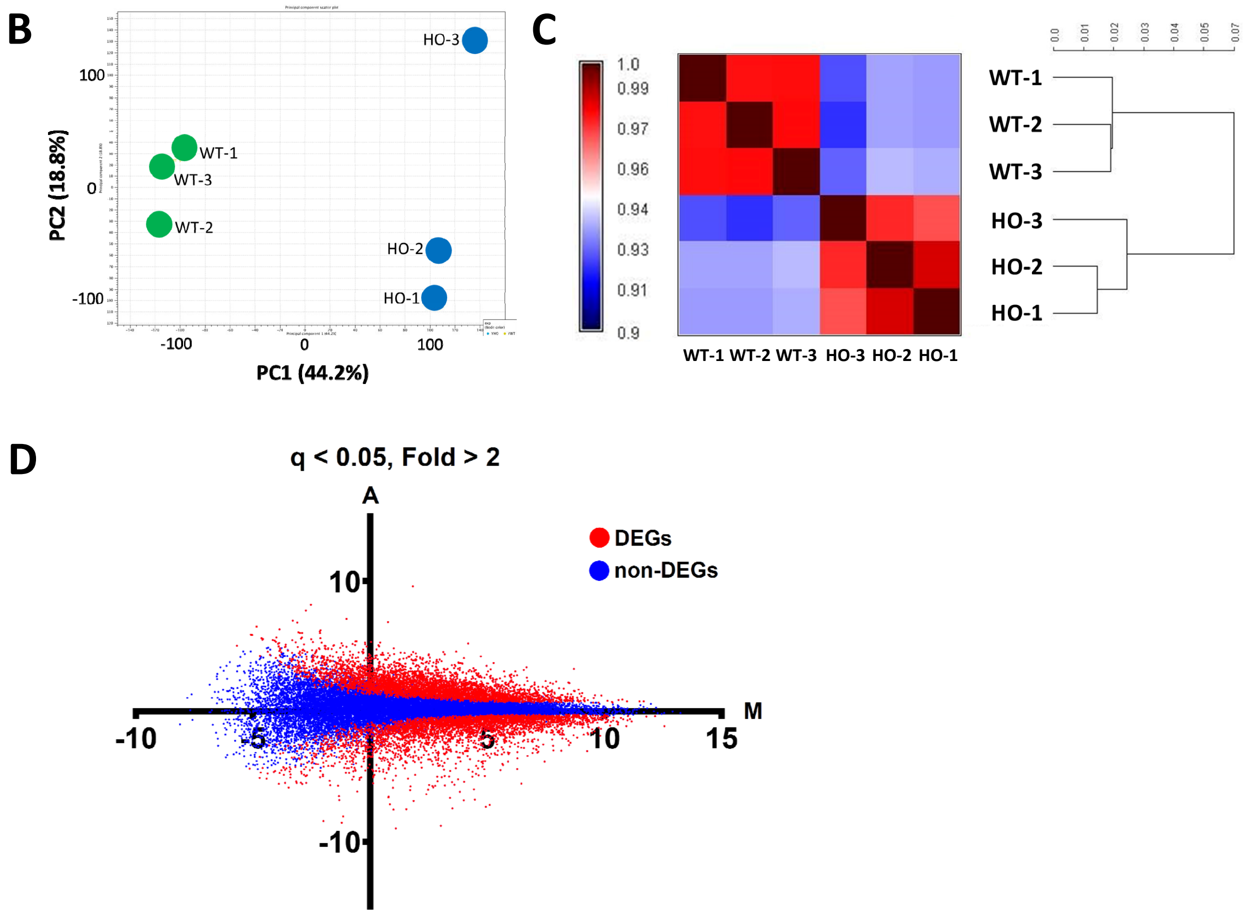


**Fig. S2. The basic analysis of RNA-seq samples. (A).** Mapping rate of reads in each sample. **(B).** Principle analysis of six independent samples showing that WT and HO samples are well separated. **(C).** Cluster analysis of six independent samples showing that WT and HO samples cluster with themselves but not with each other. **(D).** MA plot of DEGs.

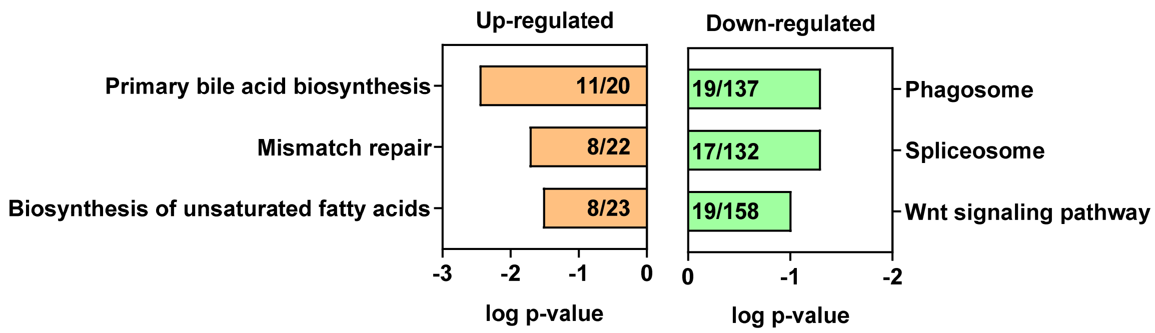


**Fig. S3. Pathway enrichment analysis of DEGs between wildtype and *foxl2l* mutant germ cells.** The numbers inside the bar indicate the ratio of DEGs to the number of all genes in this pathway.

**Fig. S4. Up-regulated DEG enriched pathways. (A).** Primary bile acid biosynthesis (dre00120). **(B).** Mismatch repair (dre03430). **(C).** Biosynthesis of unsaturated fatty acids (dre01040). DEGs in the pathways are indicated by red stars.

**A**

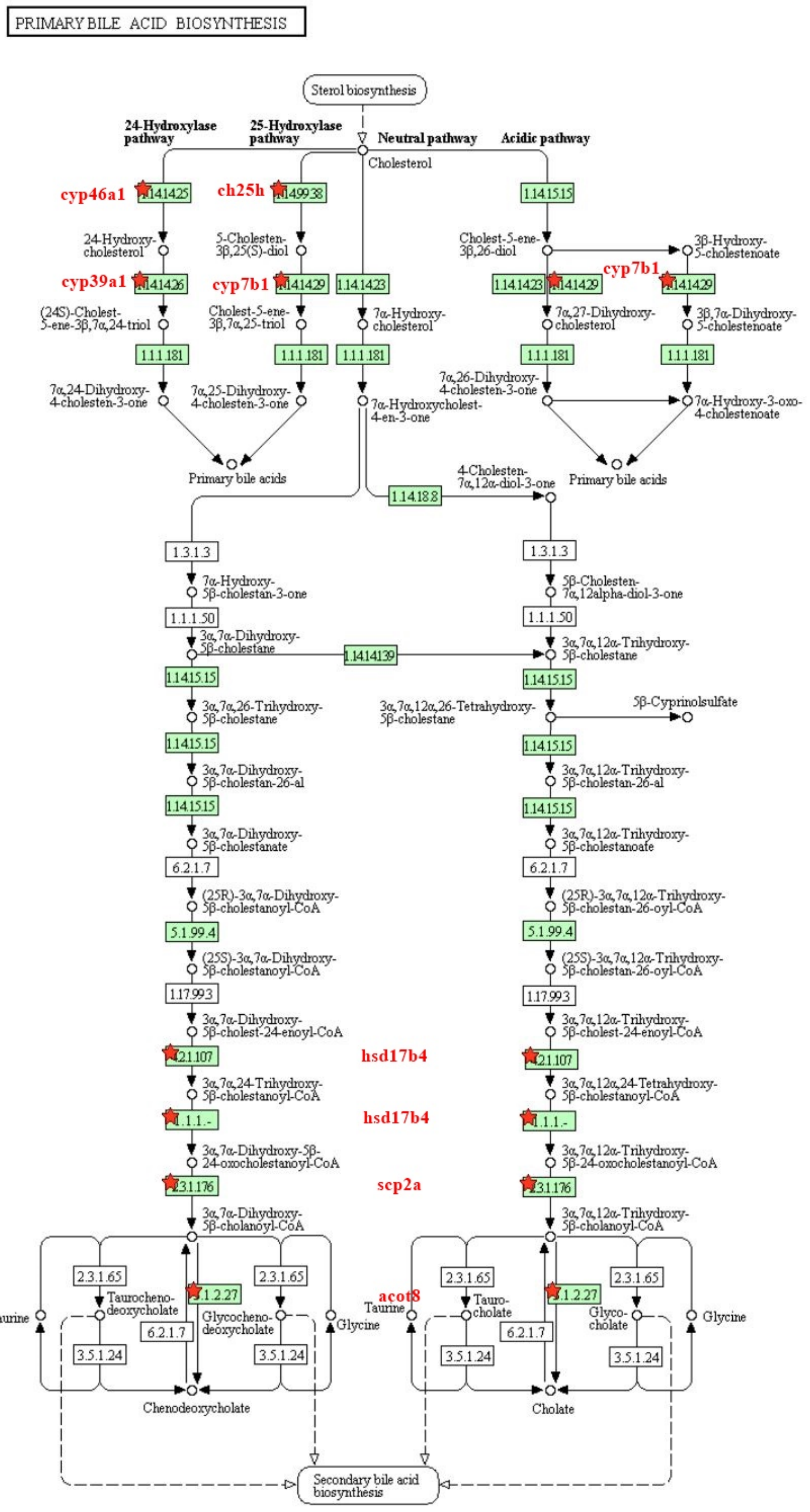

**B**

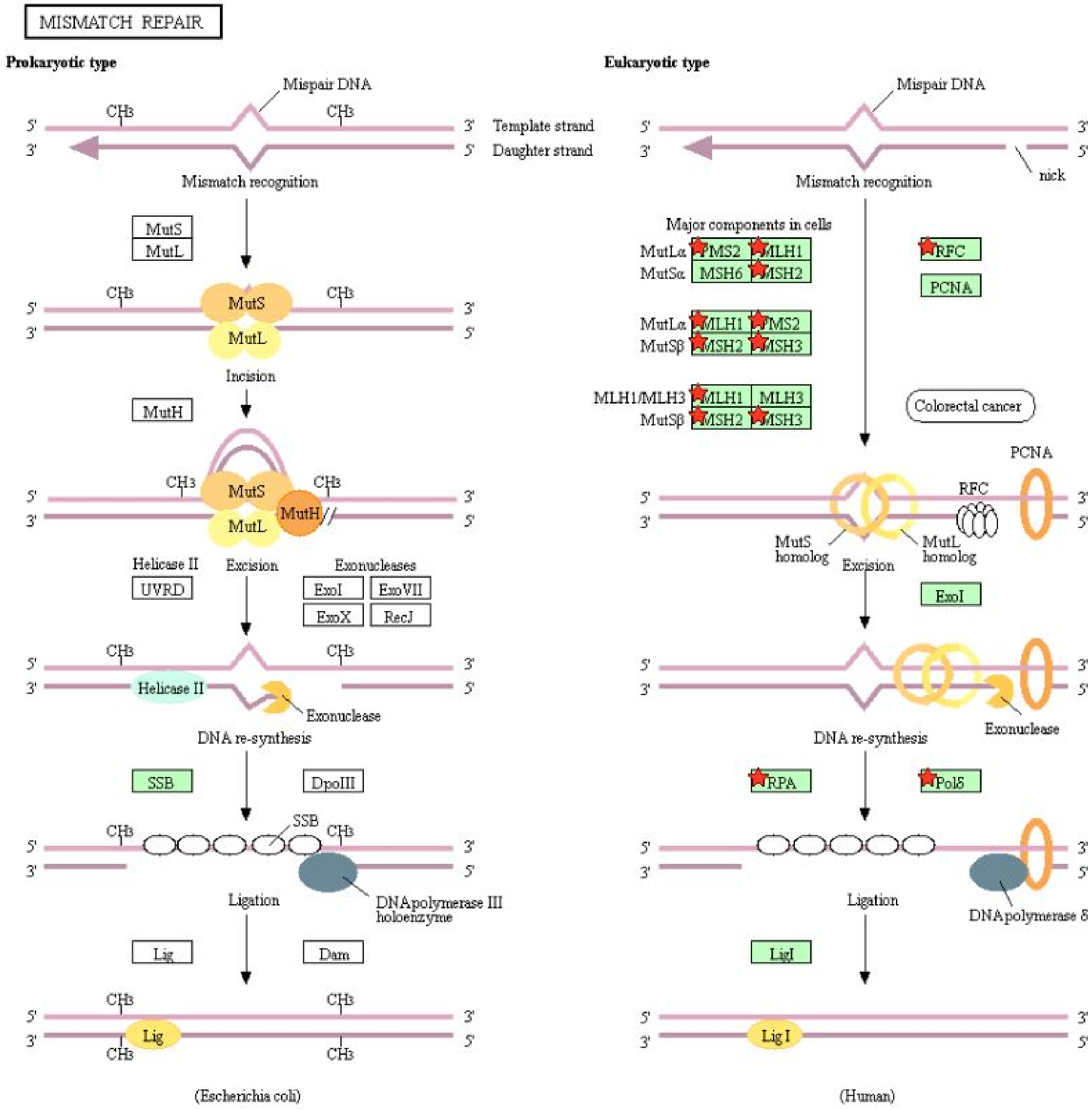
MISMATCH REPAIR
Prokaryotic type
Mispair DNA
CH3
5'
3'
Template strand
Daughter strand
Mismatch recognition
MutS
MutL
Incision
MutH
Helicase II
UVRD
Excision
Exonucleases
ExoI
ExoVII
ExoX
RecJ
Exonuclease
DNA re-synthesis
SSB
DpoIII
DNApolymerase III holoenzyme
Ligation
Lig
Dam
(Escherichia coli)
Eukaryotic type
nick
Major components in cells
MutLα PMS2 MLH1
MutSα MSH6 MSH2
MutLα MLH1 PMS2
MutSβ MSH2 MSH3
MLH1/MLH3 MLH1 MLH3
MutSβ MSH2 MSH3
RFC
PCNA
Colorectal cancer
MutS homolog
MutL homolog
ExoI
RPA
Polδ
DNApolymerase δ
LigI
Lig I
(Human)

C

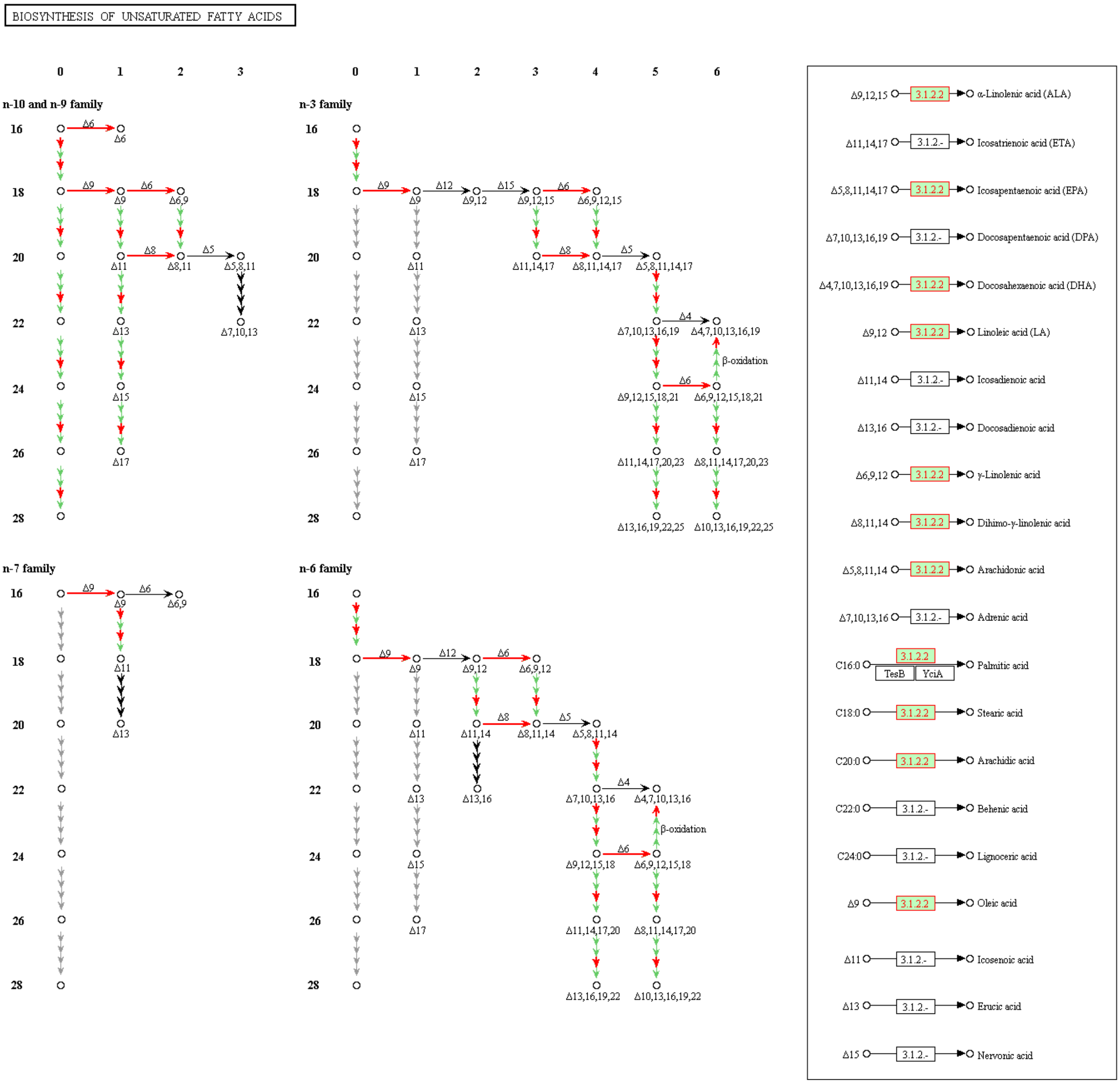

BIOSYNTHESIS OF UNSATURATED FATTY ACIDS
n-10 and n-9 family
n-3 family
n-7 family
n-6 family
β-oxidation
Δ9,12,15 3.1.2.2 α-Linolenic acid (ALA)
Δ11,14,17 3.1.2.- Icosatrienoic acid (ETA)
Δ5,8,11,14,17 3.1.2.2 Icosapentaenoic acid (EPA)
Δ7,10,13,16,19 3.1.2.- Docosapentaenoic acid (DPA)
Δ4,7,10,13,16,19 3.1.2.2 Docosahexaenoic acid (DHA)
Δ9,12 3.1.2.2 Linoleic acid (LA)
Δ11,14 3.1.2.- Icosadienoic acid
Δ13,16 3.1.2.- Docosadienoic acid
Δ6,9,12 3.1.2.2 γ-Linolenic acid
Δ8,11,14 3.1.2.2 Dihimo-γ-linolenic acid
Δ5,8,11,14 3.1.2.2 Arachidonic acid
Δ7,10,13,16 3.1.2.- Adrenic acid
C16:0 3.1.2.2 TesB YciA Palmitic acid
C18:0 3.1.2.2 Stearic acid
C20:0 3.1.2.2 Arachidic acid
C22:0 3.1.2.- Behenic acid
C24:0 3.1.2.- Lignoceric acid
Δ9 3.1.2.2 Oleic acid
Δ11 3.1.2.- Icosenoic acid
Δ13 3.1.2.- Erucic acid
Δ15 3.1.2.- Nervonic acid

**Fig. S5. Down-regulated DEG enriched pathways. (A).** Phagosome (dre04145). **(B).** Spliceosome (dre03040). **(C).** Wnt signaling pathway (dre04310). DEGs in the pathways are indicated by red stars.

**A**

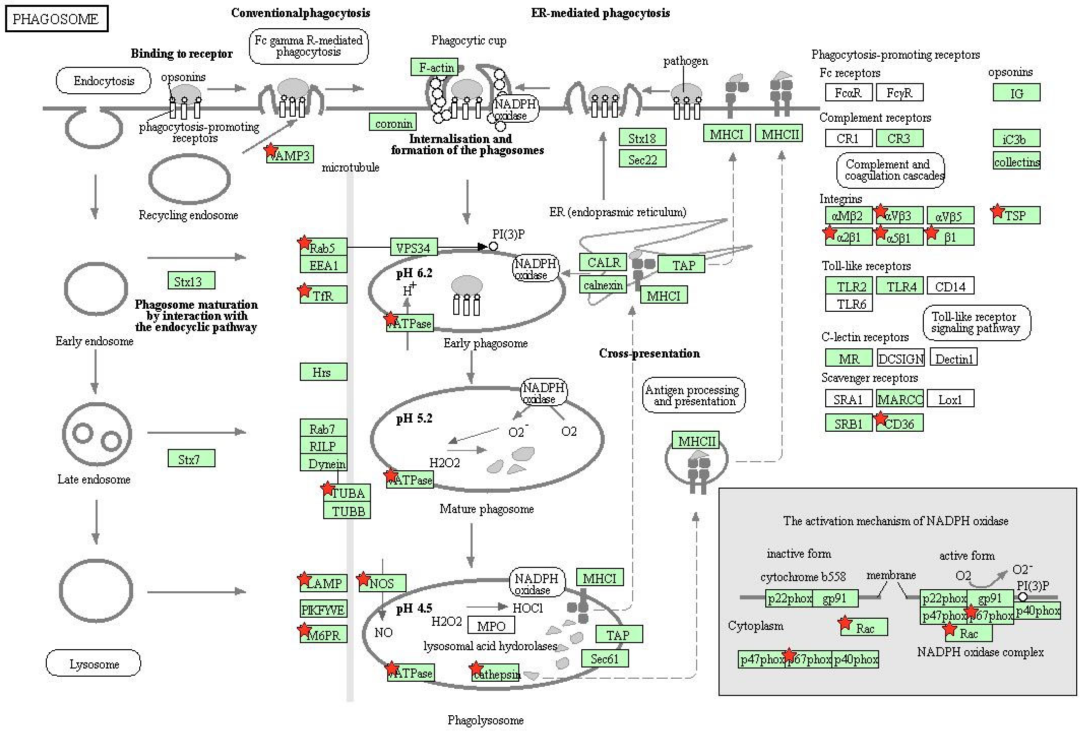


**B**

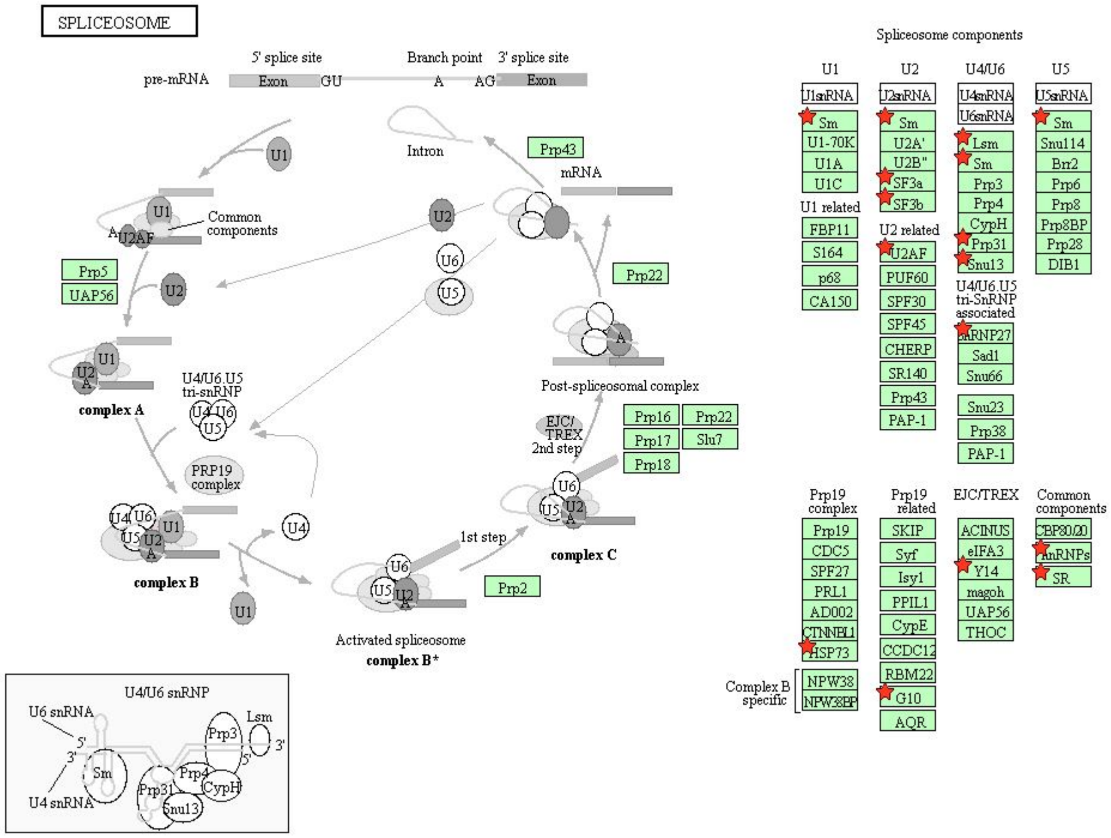

C

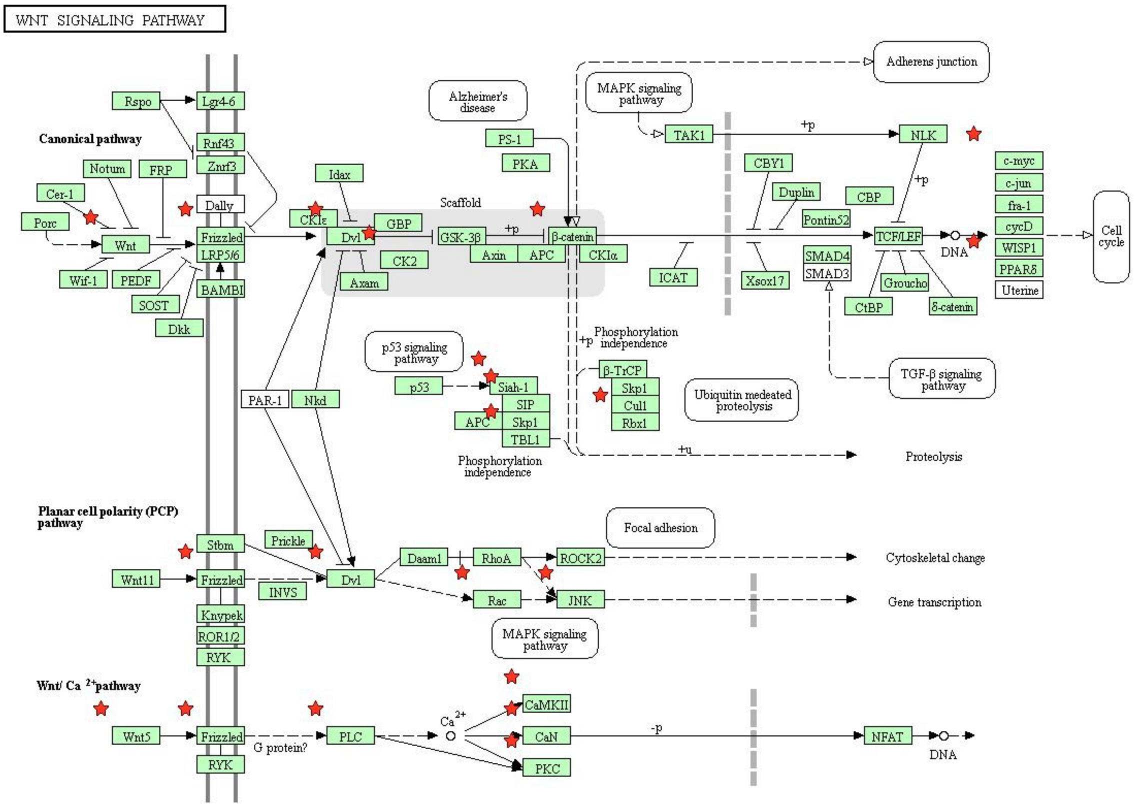
WNT SIGNALING PATHWAY
Canonical pathway
Planar cell polarity (PCP) pathway
Wnt/ Ca 2+pathway
Rspo
Lgr4-6
Rnf43
Znrf3
Notum
FRP
Cer-1
Porc
Wnt
Wif-1
PEDF
SOST
Dkk
Dally
Frizzled
LRP5/6
BAMBI
Idax
CKIε
Dvl
GBP
CK2
Axam
Scaffold
GSK-3β
Axin
APC
β-catenin
CKIα
Alzheimer's disease
PS-1
PKA
MAPK signaling pathway
TAK1
NLK
Adherens junction
CBY1
Duplin
CBP
Pontin52
TCF/LEF
DNA
ICAT
Xsox17
SMAD4
SMAD3
Groucho
CtBP
δ-catenin
TGF-β signaling pathway
c-myc
c-jun
fra-1
cycD
WISP1
PPARδ
Uterine
Cell cycle
p53 signaling pathway
p53
Siah-1
SIP
Skp1
TBL1
Phosphorylation independence
β-TrCP
Cul1
Rbx1
Ubiquitin mediated proteolysis
Proteolysis
PAR-1
Nkd
Stbm
Prickle
Wnt11
INVS
Knypek
ROR1/2
RYK
Daam1
RhoA
ROCK2
Rac
JNK
Focal adhesion
Cytoskeletal change
Gene transcription
Wnt5
G protein?
PLC
Ca2+
CaMKII
CaN
PKC
NFAT
+p
-p
+u
